# Successful spermatogonial stem cells transplantation within Pleuronectiformes: first breakthrough at inter-family level in marine fish

**DOI:** 10.1101/2021.01.29.428910

**Authors:** Li Zhou, Xueying Wang, Qinghua Liu, Jingkun Yang, Shihong Xu, Zhihao Wu, Yanfeng Wang, Feng You, Zongcheng Song, Jun Li

## Abstract

As a promising biotechnology, fish germ cell transplantation shows potentials in conservation germplasm resource, propagation of elite species, and generation of transgenic individuals. In this study, we successfully transplanted the Japanese flounder (*P. olivaceus*), summer flounder (*P. dentatus*), and turbot (*S. maximus*) spermatogonia into triploid Japanese flounder larvae, and achieved high transplantation efficiency of 100%, 75-95% and 33-50% by fluorescence tracking and molecular analysis, respectively. Eventually, donor-derived spermatozoa produced offspring by artificial insemination. We only found male and intersex chimeras in inter-family transplantations, while male and female chimeras in both intra-species and intra-genus transplantations. Moreover, the intersex chimeras could mature and produce turbot functional spermatozoa. We firstly realized inter-family transplantation in marine fish species. These results demonstrated successful spermatogonial stem cells transplantation within Pleuronectiformes, suggesting the germ cells migration, incorporation and maturation within order were conserved across a wide range of teleost species.

## Introduction

Germ stem cells have multiple differentiation potential (*Shigunov and Dallagiovanna, 2015*). They are transplanted into allogeneic or xenogeneic recipients, and then colonize, differentiate, mature and finally produce donor-derived gametes, which is called germ cell transplantation technology (GCT). GCT is a promising biotechnology for protecting endangered species as well as long-term preserving genetic resources by cryopreserved germ cells transplantation (*Lee et al., 2016; Ye et al., 2017; Yoshikawa et al., 2018*). In addition, through this technology, economic fish with long breeding cycle, large body size and low fertility can be produced by using small fish agent that is easy to raise and reproduce, which can save labor and breeding costs (*Higuchi et al., 2011; Morita et al., 2015*). It can also apply to sex control, such as producing YY supermales without administering exogenous sex steroids (*Okutsu et al., 2015*). This surrogate broodstock technology will open up a new way for fish breeding, genetic improvement, and conservation and efficient utilization of germplasm resources.

This technology originated from the 1990s, it has been greatly developed in fish in recent years (*Lin et al., 1992; Takeuchi et al., 2004; Saito et al., 2008; Wong et al., 2011; Yoshikawa et al., 2016*). Initially, GFP-labelled PGCs of rainbow trout (*O. mykiss*) were transplanted into the peritoneal cavity of the allogeneic or closely related xenogeneic species such as masu salmon (*O. masou*) at hatchling stage, resulting in production donor-derived gametes (*Takeuchi et al., 2003, 2004*). Currently, GCT has been carried out in many families, including Cyprinid (*Saito et al., 2008, 2010; Nóbrega et al., 2010*), Adrianichthyidae (*Li et al., 2016; Seki et al., 2017*), Tetraodontidae (*Hamasaki et al., 2017*), Sciaenidae (*Takeuchi et al., 2009; Yoshikawa et al., 2017*), Carangidae (*Higuchi et al., 2011; Morita et al., 2012, 2015*). Different stem germ cells including primordial germ cells (PGCs), spermatogonia, and oogonia were used as donor cells being transplanted into embryos, hatchlings and adults (*Takeuchi et al., 2004; Lacerda et al., 2013; Yoshizaki and Yazawa, 2019*). Because of the limited number of PGCs from embryos, spermatogonia were usually used as donor cells for its sexual bipotency and large amounts (*Okutsu et al., 2006, 2007; Yoshizaki et al., 2010a, b*). Therefore, spermatogonia were used for transplantation in more and more fish species instead of PGCs, especially for several marine fish species (*Takeuchi et al., 2009; Yazawa et al., 2010; Higuchi et al., 2011; Morita et al., 2012, 2015; Bar et al., 2015; Yoshikawa et al., 2017*). For example, jack mackerel (*T. japonicus*) successfully produced functional gametes of Japanese yellowtail (*S. quinqueradiata*) by intraperitoneally transplanting spermatogonia (*Morita et al., 2015*).

Recipient preparation is of prime importance for the successful transplantation of spermatogonial stem cells. Optimal recipient can greatly improve the efficiency of transplantation without immunological rejection of endogenous germ cells (*Majhi et al., 2009; Nóbrega et al., 2009; Takeuchi et al., 2020*). Sterile hybrid (*Xu et al., 2019*), triploid (*Lee et al., 2013*) or recipients with germ cell depletion induced by *dead end*-knockdown (*dnd*-MO) (*Yoshizaki et al., 2016*) or busulfan treatment (*Lacerda et al., 2010*) are commonly used for transplantation. However, not every type of sterility is easy to be obtained and applied to all species. The sterility by *dnd*-MO is just limited to model organisms, which is difficult to obtain for economic species (*Saito et al., 2008*), while busulfan treatment drastically disturbs the testicular microenvironment leading to the low efficiency, and only applies to the adult recipient producing unidirectional gamete (*Lacerda et al., 2010; de Siqueira-Silva et al., 2019*). Another type of sterility is hybrid, but whether the hybrid is sterile or not depends on the combination of the parental crosses, in which only a part of species can be obtained viable and sterile hybrids (*Xu et al., 2009; Wong et al., 2011; Piva et al., 2018*). In contrast, the triploids seem like an optimal recipient, and has been successfully used in intra- or inter-species transplantations such as rainbow trout (*Lee et al., 2013, 2016*), Grass Puffer (*T. niphoblesparents*) (*Hamasaki et al., 2017*), Niber croaker (*N. mitsukurii*) (*Yoshikawa et al., 2017*).

Turbot (*S. maximus*), Japanese flounder (*P. olivaceus*) and summer flounder (*P. dentatus*), which belong to Pleuronectiformes, are economically important marine flatfish species for aquaculture. However, since turbot and summer flounder were introduced into China, for the long-term culture and inbreeding, the germplasm resources have gradually degraded and turbot cannot spawn naturally (*Lei et al., 2012; Xu et al., 2011; Liu et al., 2015*). In previous study of flatfish, testicular germ cells of Senegalese sole (*S. senegalensis*) were transplanted into turbot larvae, and only detected proliferation in turbot juveniles, without continuous monitoring of donor cells development in the recipient (*Pacchiarini et al., 2014*). Therefore, it is necessary to explore the possibility of GCT for germplasm resources protection and surrogate broodstock among these species. In the present study, we aimed to investigate GCT within Pleuronectiformes at species, genus, and family levels. We isolated the cryopreserved spermatogonia from Japanese flounder, summer flounder, and turbot whole testes, then labeled and intraperitoneally transplanted into triploid Japanese flounder larvae, and detected the entire development process of donor-derived cells in the recipients by fluorescence tracing and molecular analysis. Finally, we successfully obtained donor-derived functional spermatozoa and offspring by artificial insemination.

## Results

### Isolation, identification and labeling of donor cells with enriched spermatogonia

In the case of turbot, the testes of sexually mature male donors contained various stages of germ cells (*Figure 1A1*). There were a large number of type A spermatogonia (Aund and Adiff) and type B spermatogonia (SpgB) distributed at the edge of the testes (*Figure 1A1’*). After whole testes cryopreservation, the isolated germ cells exhibited high survival rate (90%) by trypan blue staining (*Figure 1A1*). In order to obtain enriched spermatogonia, the isolated germ cells were subjected to Percoll density gradient centrifugation, resulting in four distinct cell bands were generated in the tube (*Figure 1A2-A5*). The top band was observed on the top of 10% Percoll, containing a high percentage spermatogonia of type A (diameter 10-15 μm) and B (diameter 6-10 μm) and a few primary spermatocytes (diameter 4-6 μm) (*Figure 1A2*); The second band on the top of 25% Percoll was thick and contained a lot of primary spermatocytes, secondary spermatocytes (diameter 3-4 μm) and a few spermatids (diameter 2-3 μm) (*Figure 1A3*); The third band was below of 35% Percoll, predominantly containing the spermatids and very few secondary spermatocytes (*Figure 1A4*); And the pellet on the bottom of the tube in 45% Percoll was the fourth band, and contained the most of spermatozoon (diameter <2 μm) and a few spermatids *(Figure 1A5*).

**Figure 1.**
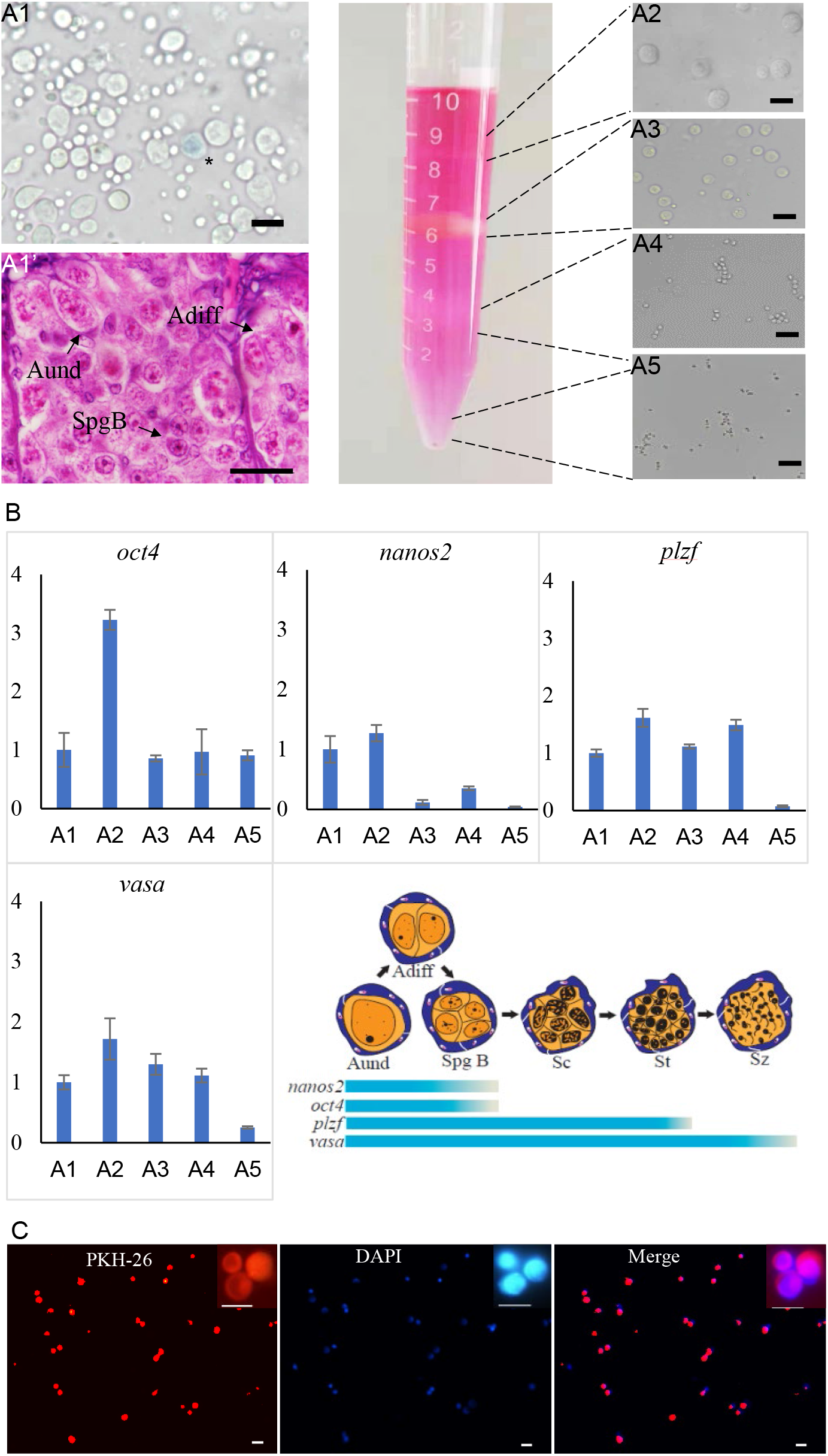
Isolation, identification and labeling of donor cells from sexually mature male turbot. (A1) The testes contained various stages of germ cells that exhibited the high rate of survival by trypan blue staining after whole cryopreservation. (A1’) Histology showed that a large number of spermatogonia (Aund, Adiff, and SpgB) distributed at the edge of the testes. (A2-A5) Four distinct cell bands were generated in the tube after Percoll density gradient (10%, 25%, 35% and 45%) centrifugation. The first cell band on the top of 10% Percoll contained abundant spermatogonia. (B) The qPCR detected the expression of *oct4*, *nanos2*, *plzf* and *vasa* genes in isolated cells. Stem-related genes *oct4*, *nanos2* and *plzf* kept higher expression in cells of the first band than others and the schematic illustration was shown below. (C) The first cell band was stained with PKH26 and DAPI. The asterisk indicated dead cell. The insets were magnification of germ cells. Aund: type A undifferentiated spermatogonia; Adiff: type A differentiated spermatogonia; SpgB: type B spermatogonia. Scale bar, 15 μm.

The qPCR detected that the expression of four genes in isolated cells (*Figure 1B*). It demonstrated that *vasa* gene well expressed in uncentrifuged cell suspension and four cell bands, however, stem-related genes *oct4*, *nanos2* and *plzf* kept higher expression in cells of the first band than others, especially for *oct4* and *nanos2* (*Figure 1B*). This identification results further confirmed that the first cell band contained ideal donor cells for transplantation.

Subsequently, the first cell band contained abundant spermatogonia was stained with PKH26 (*Figure 1C*). DAPI nuclear counterstain showed most of the cells were labeled by PKH26, of which spermatogonia accounted for 60% (*Figure 1C*). The PKH26 marker made it possible to trace donor source cells in the body cavity of recipients during early transplantation. The isolation, identification and labeling of donor cells in Japanese flounder and summer flounder were similar to those of turbot.

### Assessment the migration and colonization rates of transplanted donor cells in recipients

PKH26-labeled donor cells of turbot were observed in recipients at 14dpt, and the cells almost migrated to the genital ridge (*Figure 2B1-B3*), while the control group showed no fluorescence at the same location *(Figure 2A1-A3*). At 50 dpt, the donor cells randomly distributed and incorporated into the developing recipient gonad (*Figure 2C1-C3, D1-D3*). Like turbot, the labeled donor cells of summer flounder and Japanese flounder were also found in exposed gonads of recipients at 50 dpt, respectively (*Figure 2E1-E3, F1-F3*).

**Figure 2.**
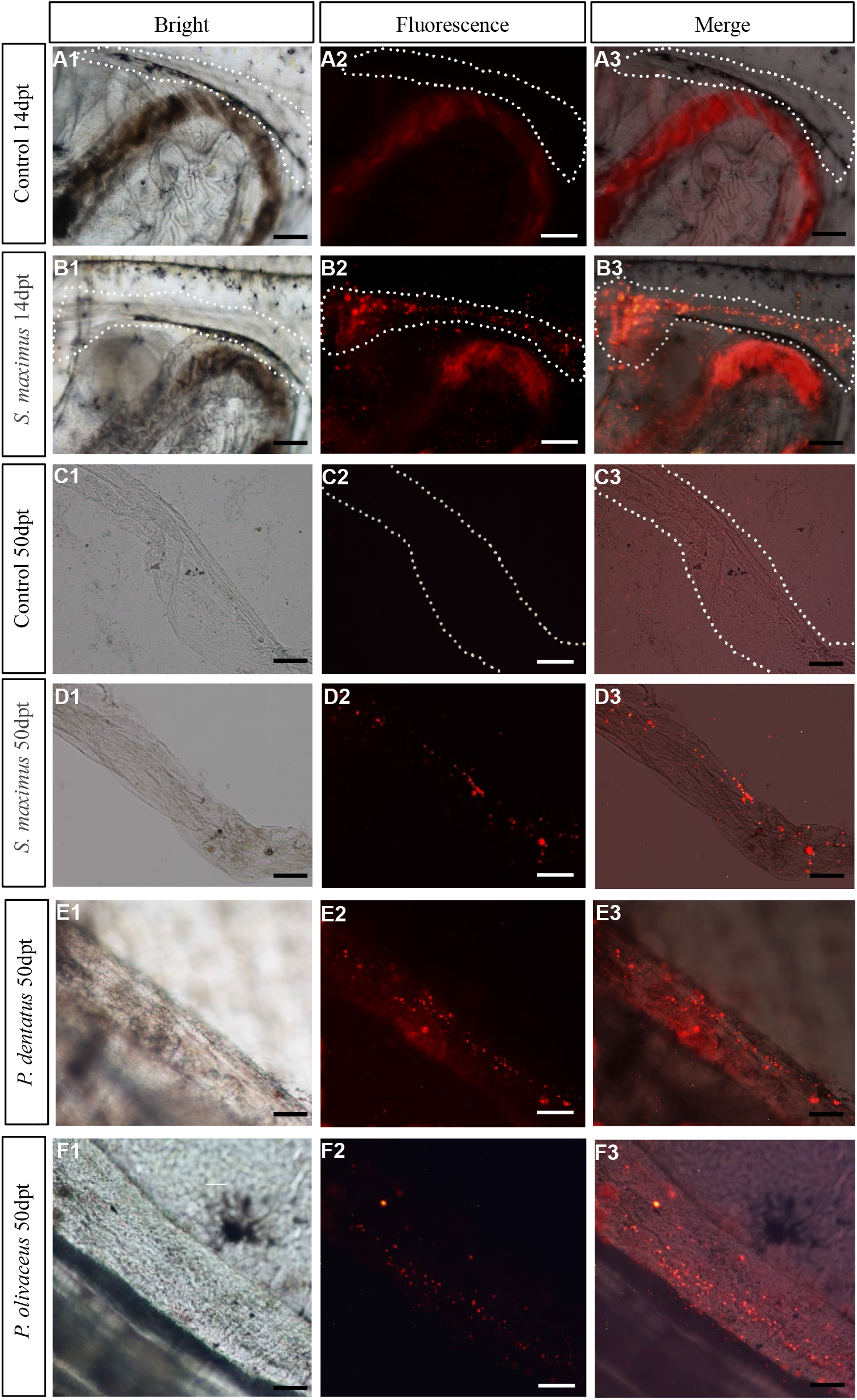
Assessment the migration and colonization rate of transplanted donor cells in recipients. (A1-A3, B1-B3) PKH26-labeled donor cells of turbot were observed at genital ridge of recipients at 14 dpt, while the non-transplanted triploid Japanese flounder controls showed no fluorescence at the same location. (C1-C3, D1-D3) At 50 dpt, the donor cells of turbot had incorporated into the developing recipient gonads, and the controls still showed no fluorescence. (E1-E3, F1-F3) The PKH26-labeled donor cells of summer flounder and Japanese flounder were also found in exposed gonads of recipients at 50 dpt, respectively. Scale bar, 100 μm.

The early assessment of transplantation efficiency by observation fluorescence in genital ridge at 14 dpt. As the days of hatching increased, the survival rate of transplanted recipients gradually increased, while the transplantation efficiency decreased (*Table 1*). And Japanese flounder (100%) and summer flounder (95.00%±0.71) had higher colonization rates than turbot (43.33%±1.53) (*Table 1*).

**Table 1.**
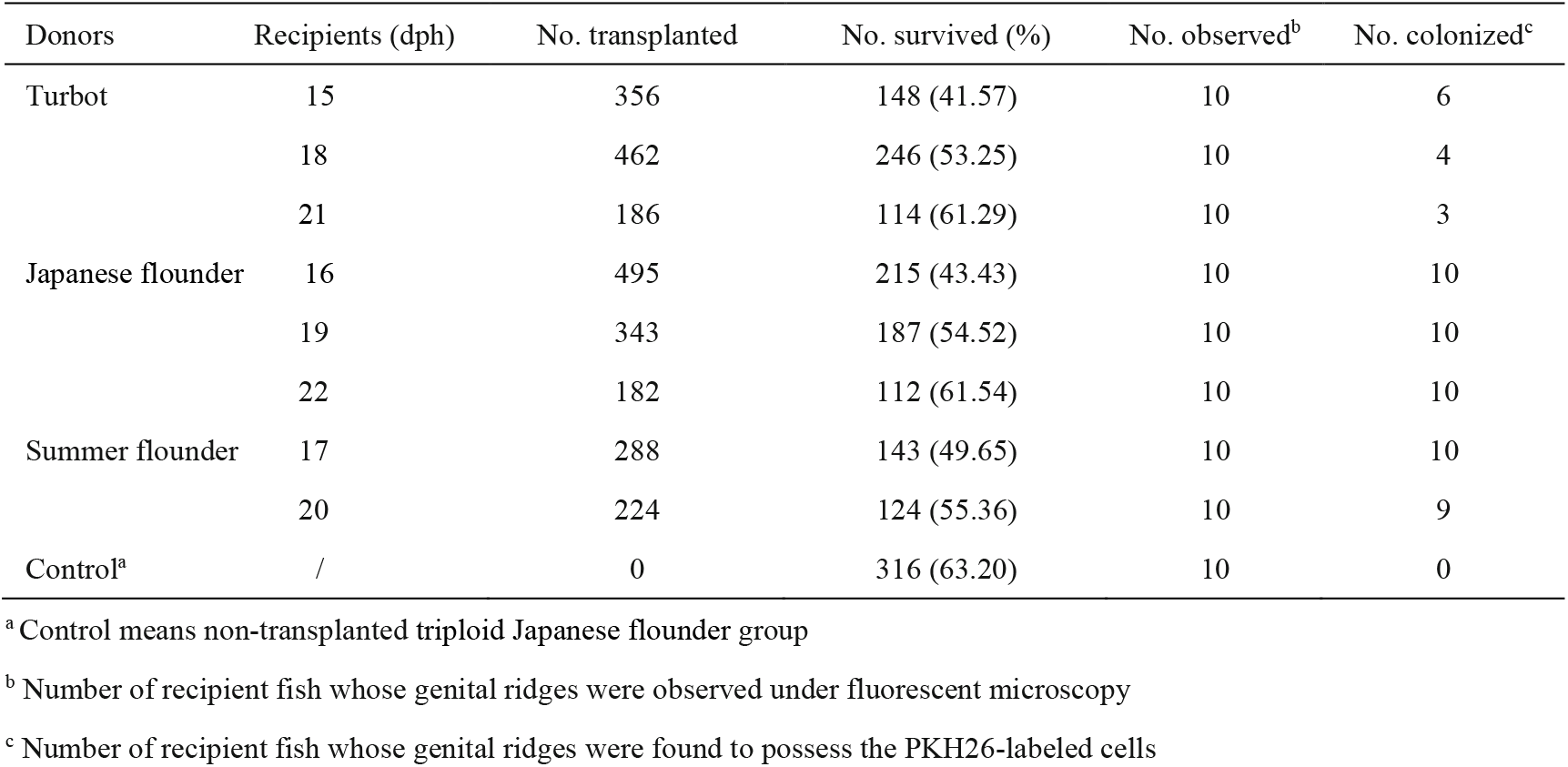
Survival of triploid Japanese flounder recipients and colonization of PKH26-positive cells in recipient genital ridges at 14 days after the allogeneic or xenogeneic transplantation

| Donors | Recipients (dph) | No. transplanted | No. survived (%) | No. observed<sup>b</sup> | No. colonized<sup>c</sup> |
| --- | --- | --- | --- | --- | --- |
| Turbot | 15 | 356 | 148 (41.57) | 10 | 6 |
|  | 18 | 462 | 246 (53.25) | 10 | 4 |
|  | 21 | 186 | 114 (61.29) | 10 | 3 |
| Japanese flounder | 16 | 495 | 215 (43.43) | 10 | 10 |
|  | 19 | 343 | 187 (54.52) | 10 | 10 |
|  | 22 | 182 | 112 (61.54) | 10 | 10 |
| Summer flounder | 17 | 288 | 143 (49.65) | 10 | 10 |
|  | 20 | 224 | 124 (55.36) | 10 | 9 |
| Control<sup>a</sup> | / | 0 | 316 (63.20) | 10 | 0 |

<sup>a</sup> Control means non-transplanted triploid Japanese flounder group
<sup>b</sup> Number of recipient fish whose genital ridges were observed under fluorescent microscopy
<sup>c</sup> Number of recipient fish whose genital ridges were found to possess the PKH26-labeled cells

### Analysis gonad development and sexual differentiation of recipients

Gonadal morphology of recipients transplanted turbot donor cells was evaluated at different development stages. Histology showed that under normal culture conditions, the ovaries of diploid turbot and Japanese flounder, including oocytes at various stages (I-IV), reached sexual maturity at 3 years old (*Figure 3A, C*). While the ovary of triploid Japanese flounder contained a large number of early oocytes (I-II) at 3 years (*Figure 3E*). The testis of diploid turbot at 2 years was found only a few spermatids and no matured spermatozoa until 3 years (*Figure 3B, B’*). However, for diploid Japanese flounder, the spermatozoa matured at 2 years old (*Figure 3D*). Although the matured triploid Japanese flounder contained spermatozoa, most of them were deformed and lacked flagellum (*Figure 3F*). For the types of spermatozoa maturation, in Japanese flounder, the germ cells at various stages developed in the spermatocysts arranged at the periphery of the lobules till the spermatozoa, then released to the lobular Lumen (*Figure 3B*). However, in turbot, when the germ cells developed to the secondary spermatocytes stage, the boundary between the spermatocysts became not obvious, and mixed together (*Figure 3D*).

**Figure 3.**
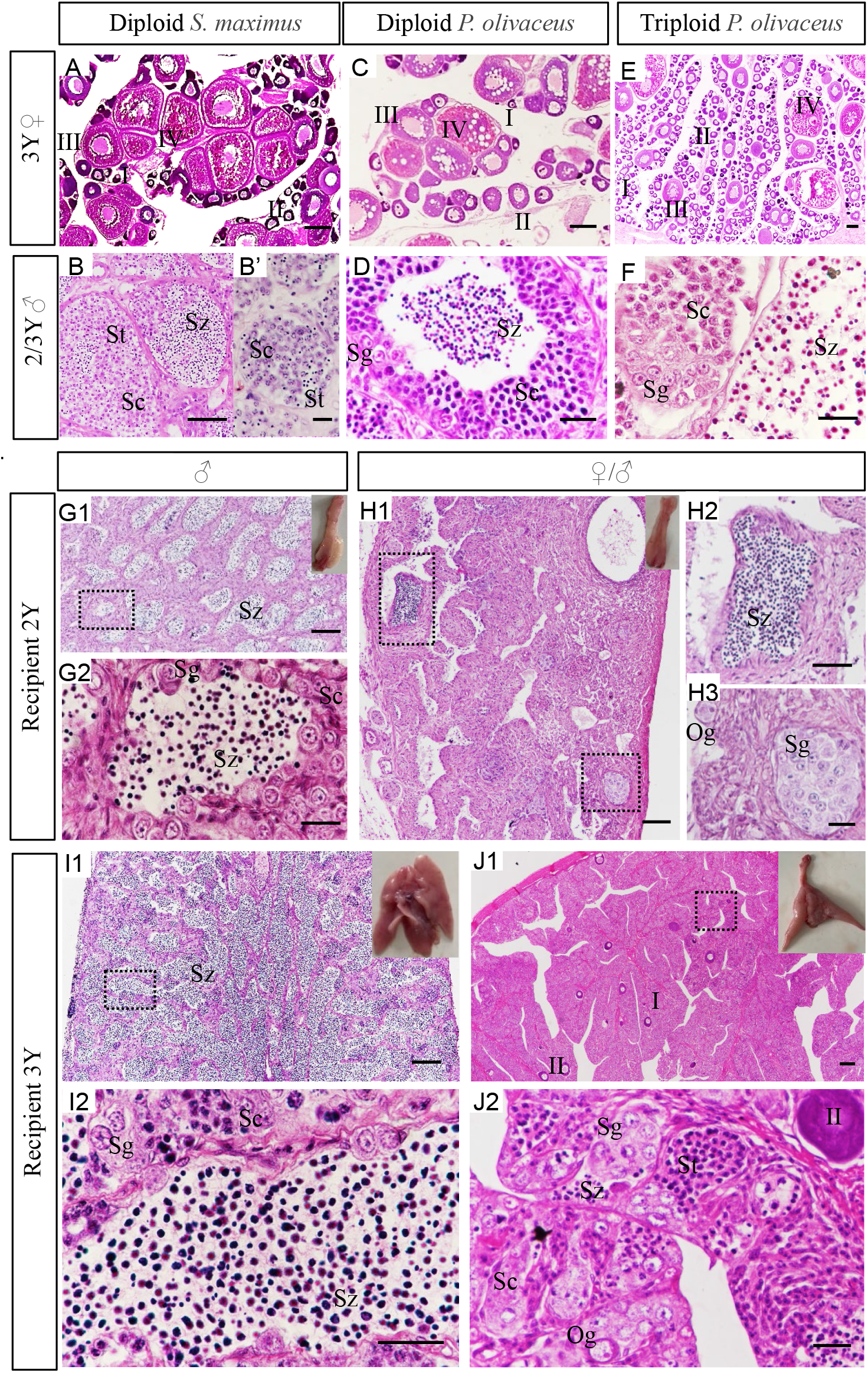
Analysis gonad development and sexual differentiation of recipients by histology. (A-F) The gonads of controls at 2-3 years old. (A, C, E) The ovaries of diploid turbot and Japanese flounder at 3years old included oocytes at various stages (I-IV), while triploid Japanese flounder contained a large number of early oocytes (I-II). (B-B’) The testis of diploid turbot at 2 years was found only a few spermatids and no matured spermatozoa until 3 years. (D) However, for diploid Japanese flounder, the spermatozoa matured at 2 years old. (F) The matured triploid Japanese flounder contained deformed spermatozoa. (G1-J2) The gonads of recipients transplanted turbot spermatogonia at 2-3 years old. (G1-G2, I1-I2) The appearance of recipient gonads was identified as testes, which showed typical structure of lobular-type. The testis contained germ cells of various stages at 2ypt and produced a large number of spermatozoa at 3 ypt. (H1-H3, J1-J2) The appearance of recipient gonads was identified as ovaries, which showed the structural characteristics of the ovigerous lamella. But besides oogonia, it also contained a few spermatids even spermatozoa at 2 ypt. When they developed for 3 years old, a few early oocytes (I-II) were found, and a number of male germ cells at various stages were also observed. G2, H2, H3, I2, J2 were the amplification of black dotted frame in G1, H1, I1, J1, respectively. The insets showed appearance of chimeric gonads. Sg, spermatogonia; Sc1, primary spermatocytes; Sc2, secondary spermatocytes; St, spermatids; Sz, spermatozoon. Scale bar, 100 μm (A, C, E, G1, H1, I1, J1); 50 μm (B, D, F, G2, H2, H3, I2, J2); 10 μm (B’).

Generally, the gonads of flatfish are paired, the testes are bilobed structure, and the ovaries are club-shaped. The appearance of recipient gonads was identified as testes, which showed typical structure of lobular-type on tissue sections (*Figure 3G1-G2, I1-I2*). The testis was composed of many spermatocysts, including male germ cells at different developmental stages, such as spermatogonia and spermatocytes, spermatids and spermatozoa in recipients at 2ypt (*Figure 3G1-G2*). The testis of recipient at 3ypt was more matured and produced a large number of spermatozoa (*Figure 3I1-I2*).

According to the appearance, the gonads of recipients at 2ypt were identified as ovaries, which showed the structural characteristics of the ovigerous lamella on tissue sections (*Figure 3H1-H3*). But besides oogonia, it also contained a few spermatids even spermatozoa (*Figure 3H1-H3*). When they developed for 3 years old, a few early oocytes (I-II) were found, and a number of male germ cells at various stages were also observed, containing spermatogonia, spermatocytes, and even some spermatids and spermatozoa (*Figure 3J1-J2*). There were two different types of germ cells in recipient gonads, including male and female germ cells, which were referred to intersex (*Figure3 H1-H3, J1-J2*). Sex ratio was calculated according to the anatomy and histology of the gonads. In short, the ratio of bisexual to male was about 1:1, as shown in *Table 2*.

**Table 2.**
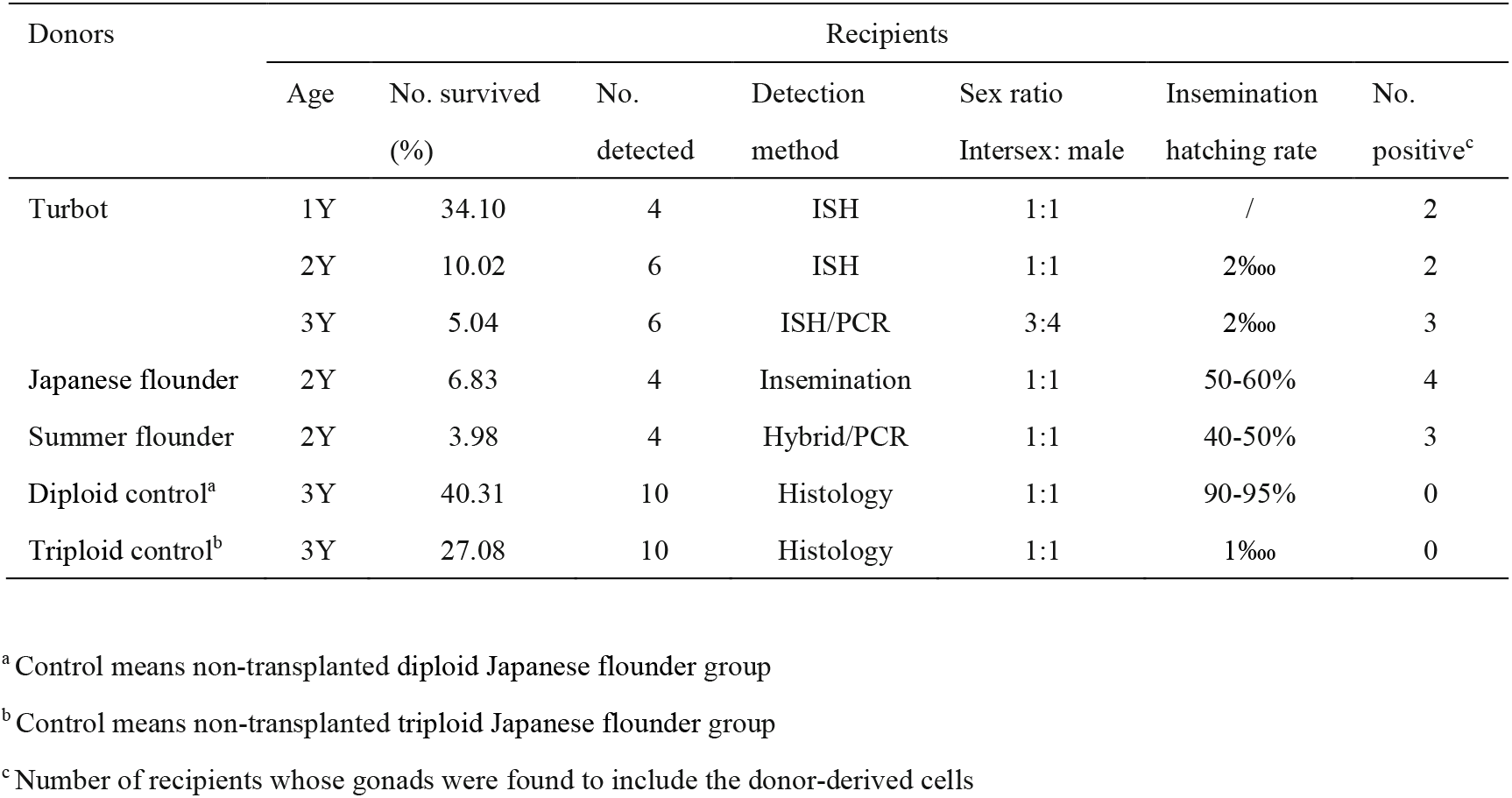
Detection donor-derived cells from recipients at different stages

### Detection donor-derived cells and proliferation in recipients

The *vasa* gene was used as a specific marker of germ cells for identifying turbot germ cells from transplanted triploid Japanese flounder gonad. ISH confirmed that both probes were specifically hybridized with *vasa* mRNAs from turbot and Japanese flounder gonads, respectively (*Figure 4A1-A8*). And the *vasa* signals were predominantly detected in early germ cells, such as spermatogonia, spermatocytes, oogonia and early oocytes (I-III), and no signal was detected in spermatids, spermatozoon, and relatively mature oocytes (*Figure 4A1-A8*). When serial gonad sections of 125dpt recipient were hybridized with species-specific probes, the Japanese flounder *vasa* signals were widely presented in the whole gonadal germ cells, while the turbot signals only existed in a cluster of germ cells (*Figure 4B1-B2*). Refer to H&E staining, the germ cells were mostly composed of SpgB (*Figure 4B3*).

**Figure 4.**
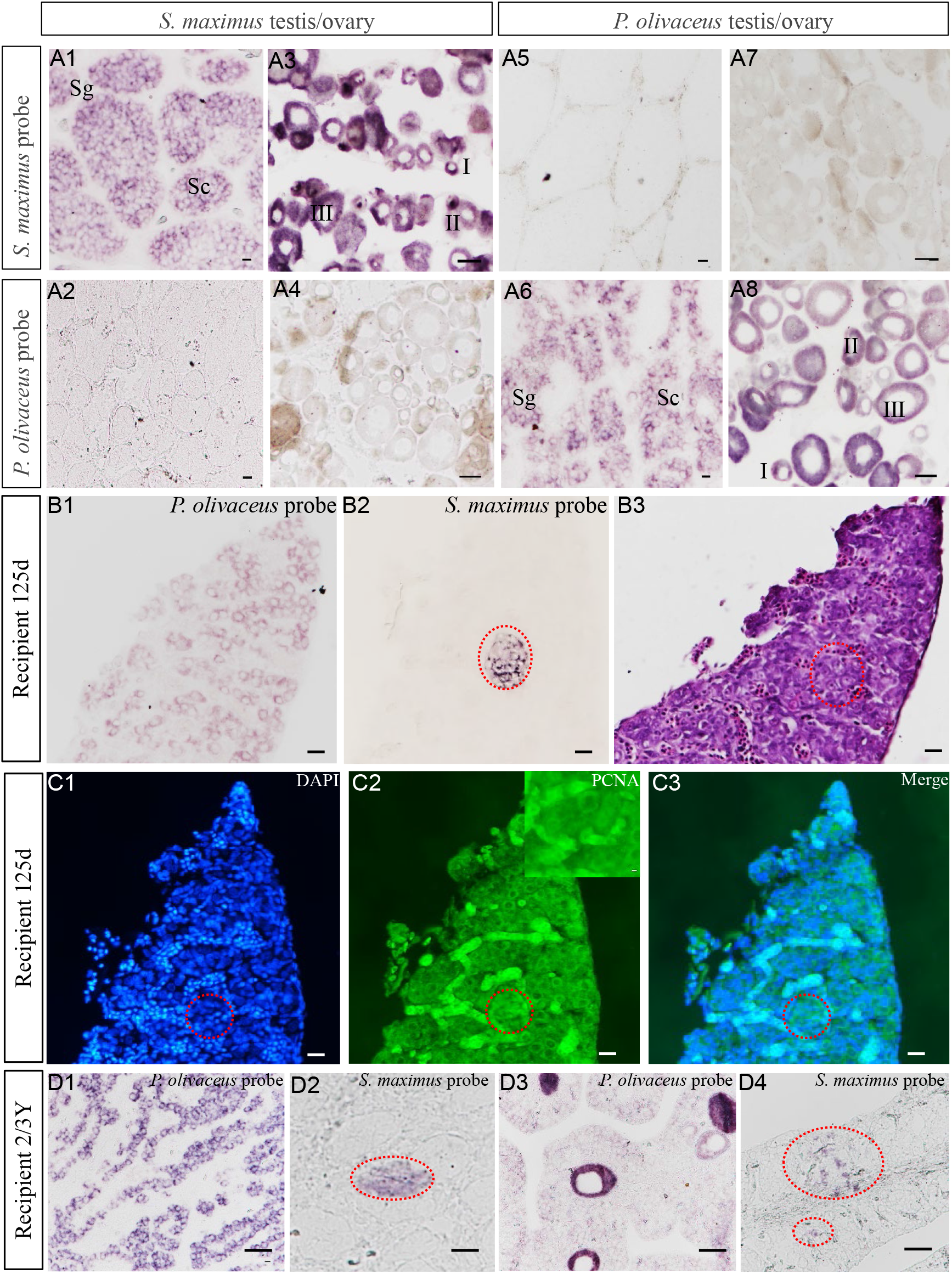
Detection donor-derived cells and proliferation in recipients by ISH and immunohistochemistry. (A1-A8) ISH confirmed that both probes were specifically hybridized with *vasa* mRNAs from turbot and Japanese flounder gonads, respectively. (B1-B3) Detected turbot cells from triploid Japanese flounder recipients at 125dpt by ISH with species-specific probes. The Japanese flounder *vasa* signals were widely presented in the germ cells of recipient, while the turbot signals only existed in a cluster of germ cells. Refer to H&E staining, the germ cells were mostly composed of SpgB. (C1-C3) Detected donor-derived cells proliferation by immunohistochemistry with anti-PCNA antibody. The section was treated with DAPI and anti-PCNA antibody after ISH, and merged. The germ cells of turbot proliferated in triploid Japanese flounder gonad. (D1-D2) Japanese founder and turbot male germ cells of spermatogonia and spermatocytes were identified from the gonad of male chimeras at 2yt, respectively. (D3-D4) Turbot male germ cells were also identified in intersexual chimeras at 3yt, which also included Japanese founder early oocyte and male germ cells. The red dotted circle represented the turbot signal area of ISH, and the inset was amplified to the signal area. Scale bar, 10 μm (A1, A2, A5, A6, B1-C3); 50μm (A3, A4, A7, A8, D1-D4).

Cell proliferation was detected at 125dpt recipient by immunohistochemistry with anti-PCNA antibody after ISH. The SpgB were identified by DAPI staining and showed strong green fluorescence after treated with anti-PCNA antibody (*Figure 4C1-C2*). The merged image demonstrated the proliferation of germ cells, including the donor-derived cells (*Figure 4C3*). These results showed that the germ cells of turbot proliferated in the somatic microenvironment of triploid Japanese flounder gonad.

Similarly, at 2-3 years old, the donor-derived germ cells, such as spermatogonia and spermatocytes, were identified from transplanted triploid Japanese flounder gonads (*Figure 4D1-D4*). Interestingly, we found turbot germ cells of spermatogonia and spermatocytes from the intersexual recipient gonad, which also included Japanese founder early oocyte and male germ cells (*Figure 4D3-D4*). All of the above results showed that donor-derived cells existed throughout the development of the recipient gonads

### Identification donor-derived spermatozoa or offspring from matured recipients

The morphology and function of donor-derived spermatozoa from inter-family transplantation were identified by scanning electron microscopy, PCR and artificial insemination. The spermatozoa of both species consisted of three parts: head, mid-piece and flagellum (*Figure 5A1-A4*). However, the shape of spermatozoa head was different. For Japanese flounder spermatozoa, the long diameter was 1.60±0.50μm, and the short diameter was 1.50±0.50μm (*Figure 5A1-A2*). While for turbot, due to the anterior pit of nucleus, the long diameter was 1.70±0.50μm, and the short diameter was 0.80±0.50μm (*Figure 5A3-A4*). The spermatozoa of 5 recipients at 2 ypt were observed, in which the recipients 3# and 5# contained scattered turbot spermatozoa with anterior pit of nucleus (*Figure 5A5-A6*). Besides, the recipients contained a large number of triploid Japanese flounder spermatozoa with malformed or totally absent from flagellum (*Figure 5A5-A6*). Then, PCR was performed with species-specific primers to identify turbot DNA from the spermatozoa in 5 recipients. The results showed that all 5 recipients contained Japanese flounder DNA (*Figure 5B1*). And the turbot DNA was also detected in the recipients 3# and 5# by nested PCR (*Figure 5B1’*). In order to further analyze the spermatozoa function, artificial insemination was carried out for 5 recipients. The larva of turbot and the Japanese flounder showed different body colors. Turbot larvae showed obvious orange pigment clusters from somite stage on the dorsal and anal fin folds, while Japanese flounder larvae was covered with melanin dots (*Figure 5C1-C3*). The fertilization rate and hatching rate of the five recipients were kept at a very low level (2%oo), only the recipients 3# and 5# could produce 2 turbot-like offspring, respectively (*Table 2, Figure 5C4*). Because the hybrid between turbot and Japanese flounder could not develop normally, the 2 offspring obtained from artificial insemination were donor derived.

**Figure 5.**
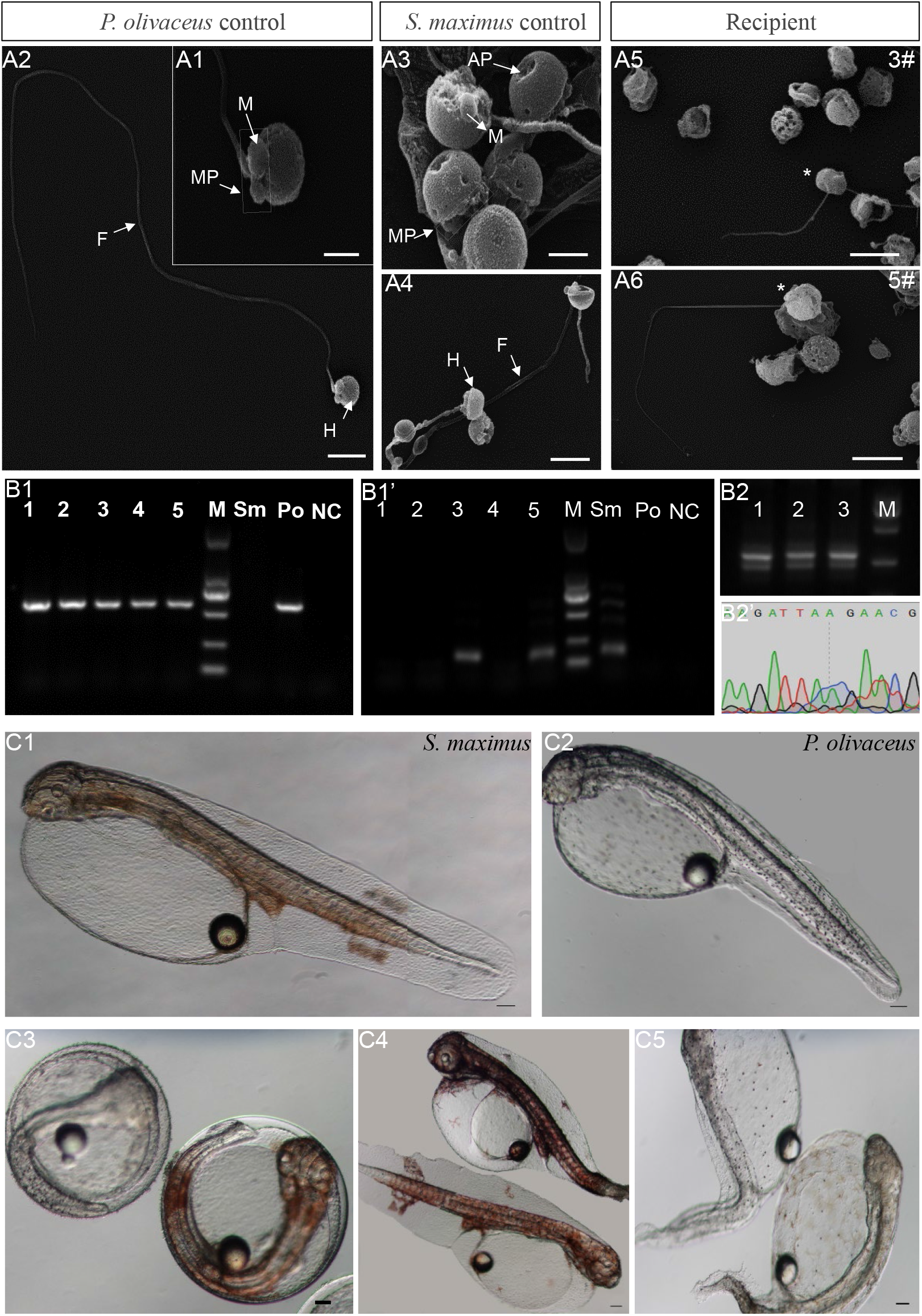
Identification donor-derived spermatozoa or offspring from matured recipients. (A1-A6) Turbot spermatozoa morphology was identified from inter-family transplantation by scanning electron microscopy. The recipients 3# and 5# contained scattered turbot spermatozoa with anterior pit of nucleus. (B1-B1’) Identify turbot DNA of the spermatozoa from 5 recipients by nested PCR. All 5 recipients contained Japanese flounder DNA, and only the recipients 3# and 5# also contained turbot DNA. (B2-B2’) The hybrids from 3 recipients showed that there were two bands with similar molecular sizes on 2% agarose gel electrophoresis. After sequencing, double peaks appeared in the different sequence regions of Japanese flounder and summer flounder. (C1-C3) The larva of turbot and the Japanese flounder showed different body colors. Turbot larvae showed obvious orange pigment clusters from somite stage on the dorsal and anal fin folds, while Japanese flounder larvae was covered with melanin dots. (C4) The spermatozoa function was identified by artificial insemination, and only the recipients 3# and 5# could produce 2 turbot offspring, respectively. (C5) As control, the embryos from triploid were all malformed and died during the process of development. The asterisk indicated turbot spermatozoa. H: head; MP: mid-piece; F: flagellum; M: mitochondrion; AP: Anterior pit of nucleus. Scale bar, 1 μm (A1, A3); 2.5μm (A2, A4); 4μm (A5, A6); 100μm (C1-C5).

And the triploid Japanese flounders were transplanted with allogeneic spermatogonia, the fertilization rate was as high as 90%. But some embryos died at the blastula stage, and others gradually stabilized after the gastrulation, resulting in the hatching rate at 50-60% (*Table 2*). As controls, the embryos from diploid developed well under breeding conditions, keeping the fertilization rate and hatching rate at high level (90-95%), and the embryos from triploid were all malformed and died during the process of development (*Table 2, Figure 5C5*).

At the same time, the spermatozoa of 3 recipients transplanted summer flounder germ cells were hybridized with Japanese flounder eggs. The hatching rate of recipients 1# and 3# were 50-60%, which was not significantly different from above intra-species transplantation (*Table 2*). And the recipients 2# kept 30%, which was relatively low (*Table 2*). The PCR results of hybrids from 3 recipients showed that there were two bands with similar molecular sizes on 2% agarose gel electrophoresis. After sequencing, double peaks appeared in the different sequence regions of Japanese flounder and summer flounder, which further confirmed that they contained the DNAs of two species (*Figure 5B2-B2’*).

## Discussion

The present study demonstrated the successful intra-species, intra-genus and inter-family GCT in Pleuronectiformes. We transplanted Japanese flounder, summer flounder, and turbot spermatogonia into the peritoneal cavities of triploid Japanese flounder larvae, and achieved donor-derived functional spermatozoa from recipients, respectively (*Figure 6*). This firstly realized inter-family transplantation in marine fish species and shortened the maturation time of turbot spermatozoa.

**Figure 6.**
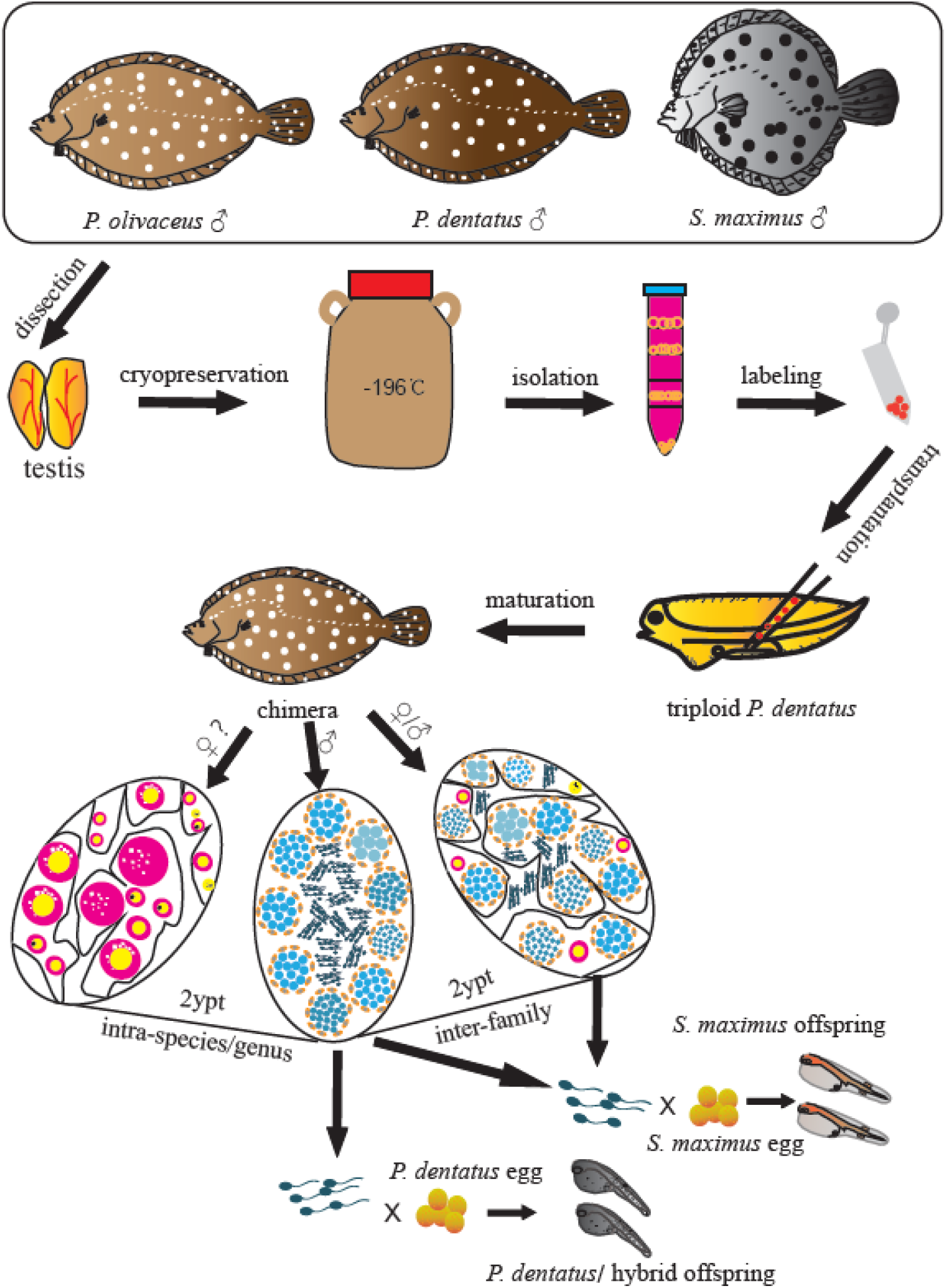
Schematic illustration of spermatogonial stem cell transplantation whin Pleuronectiformes. The spermatogonia of Japanese flounder, summer flounder, and turbot were isolated from cryopreserved whole testes, then labeled and transplanted into the peritoneal cavities of triploid Japanese flounder larvae. The transplanted recipients differentiated into male and female chimeras in intra-species and intra-genus transplantations, and only male and intersex chimeras in inter-family transplantations. Finally, donor-derived spermatozoa and the offspring by artificial insemination were obtained from recipients at 2ypt, respectively.

In previous studies, fish GCT within family level had been reported in many species, including transplanting Niber croaker spermatogonia into allogeneic larvae (intra-species, *Yoshikawa et al., 2016*), Pejerrey (*O. bonariensis*) spermatogonia into adult patagonian pejerrey (*O. hatcheri*) (intra-genus, *Majhi et al., 2014*), and Atlantic salmon (*S. salar*) spermatogonial cells into rainbow trout larvae (inter-genus, *Hattori et al., 2019*). And donor-derived gametes and offspring were successfully produced. However, as the phylogenetic distance between the donor and recipient gradually becomes greater, especially for species belonging to a different family or order, the transplantations are difficult to succeed. So far, in freshwater fish, only the successful inter-family transplantation of loach and zebrafish and inter-order transplantation of Jundia catfish (*R. quelen*) and Nile tilapia has been reported (*Saito et al., 2008; Silva et al., 2016*). In marine fish, there is no successful transplantation beyond family level. Due to the large body size, long maturation period, complicated reproduction regulation and cultivation conditions of commercially important species, the transplanted germ cells often cannot develop and mature in the recipients (*Yazawa et al., 2010; Higuchi et al., 2011; Bar et al., 2015*). The inter-family transplantations were tried in southern bluefin tuna (*T. maccoyii*) to yellowtail kingfish (*S. lalandi*), as well as Japanese yellowtail to Niber croaker, the transplanted cells only migrated to the genital ridge and proliferated in developing gonad, but disappeared at later stages (*Higuchi et al., 2011; Bar et al., 2015*). In this study, we successfully transplanted the Japanese flounder, summer flounder, and turbot spermatogonia into triploid Japanese flounder larvae. Transplantation efficiency of turbot-Japanese flounder (33-50%), summer flounder-Japanese flounder (75-95%) and Japanese flounder-Japanese flounder (100%) was achieved by fluorescence tracking and molecular analysis. Subsequently, the donor-derived spermatozoa function was further confirmed by artificial insemination. These results demonstrated germ cells from allogeneic or xenogeneic species within Pleuronectiformes were able to migrate, colonize, proliferate, differentiate and mature in recipients, and finally produce donor-derived spermatozoa and offspring. This also indicated even in divergent phylogenetic relationship, donor-derived germ cells could recognize and respond to the guide signals under the regulation of the recipient developmental environment, eventually mature in the recipient (*Saito et al., 2008; Silva et al., 2016*).

Interestingly, we also found that the sex differentiation of recipients was influenced by donor germ cells. The previous study showed that recipients in the transplantations between species of closed phylogenetic relationship could produce functional donor-derived spermatozoa and eggs (*Takeuchi et al., 2004; Yoshizaki et al., 2016; Hamasaki et al., 2017; Farlora et al., 2018*). However, transplantations between species of highly divergent in evolution, the recipients usually differentiated into males, but only a few of them produced functional spermatozoa (*Saito et al., 2008; Morita et al., 2015; Silva et al., 2016; Shang et al., 2018*). Donor germline stem cells produce gametes based on the gender of the recipient, not their own gender (*Yoshizaki et al. 2010b*). In this study, we obtained normal developed male and female chimeras in intra-species and intra-genus transplantations, and the male chimeras produced the functional spermatozoa, although the female chimeras were currently immature. In inter-family transplantation, we only obtained male and intersex chimeras, with a ratio of approximately 1:1, and no female chimeras. The gonad of male chimeras also developed normally, and produced donor-derived functional spermatozoa. The gonad of intersex chimeras showed ovary morphology and structure, however it contained turbot male germ cells that could continue to develop into functional spermatozoa, besides Japanese flounder female (a few early oocytes) and male germ cells. Compared with the non-transplantation female triploid Japanese flounder control, the introduction of exogenous spermatogonia inhibited the development of oocytes in the recipients, and promoted the regulation of the recipient to develop toward masculinization. That indicated that the recipients of Japanese flounder sex differentiation direction were more easily to be influenced by external germ stem cells.

In addition, Japanese flounder produced turbot spermatozoa, which shortened the turbot spermatozoa production time from 3 year to 2 year. Under artificial breeding conditions, turbot generally does not reach sexual maturity until three years old. And turbot spermatogenesis type belongs to semicystic type, in which the cyst ruptures at the spermatocyte or spermatid stage, so spermatozoa mixed other germ cells are released into the lumen (*Sàbat et al., 2009; Mazzoldi, 2001*). In contrast, the Japanese flounder belongs to cystic type, in which the spermatogenesis process takes place entirely inside the cyst and the spermatozoa are released into the lumen after the cyst breaks (*Biagi et al., 2016*). The histology results showed the turbot spermatogenesis conformed to the recipients during the process of testis development and maturation. In inter-family transplantation between Japanese flounder and turbot, the Japanese flounder still could surrogate the turbot for gametes production, although the donor and recipient required different water temperature and photoperiod control to regulate gonad maturation, even their spermatogenesis belonged totally two different spermatogenesis types, suggesting that the mechanisms underlying donor germ cells migration, incorporation and maturation in recipients were conserved across a wide range of teleost species (*Takeuchi et al., 2004; Saito et al., 2008; Silva et al., 2016; Hattori et al., 2019*). Moreover, the turbot cells were transplanted at hatched Japanese flounder larvae stage, and remained in the recipients for a longer period (more than 2 years) compared with previous distant transplantations (*Saito et al., 2008; Higuchi et al., 2011; Bar et al., 2015; Silva et al., 2016*). It is totally possible to transplant germ cells from large-bodied commercially important marine fish species and endangered species to related small and easier to breed species.

In conclusion, we successfully transplanted the Japanese flounder, summer flounder, and turbot spermatogonia into triploid Japanese flounder larvae, and achieved donor-derived functional spermatozoa from recipients, respectively. We also found that the transplanted recipients differentiated into male and female chimeras in intra-species and intra-genus transplantations, and only male and intersex chimeras in inter-family transplantations. Moreover, the intersex chimeras could mature and produce turbot functional spermatozoa. This firstly realized inter-family transplantation in marine fish species and shortened the maturation time of turbot spermatozoa. The results of this study promote the understanding the mechanism of the germ stem cells differentiation and maturation, and provide an alternative approach for the breeding of marine fish and the preservation of genetic resources.

## Materials and methods

### Experimental animals and general rearing protocols

In this study, all the fish were reared at Shenghang Sci-Tech Co., Ltd. (Weihai, Shandong Province, China). For donor preparation, sexually mature male individuals were cultured in 30m^3^ circular tanks with flowing seawater, and fed with frozen fish. For recipient preparation, the Japanese flounder broodstock was raised in a circular tank, and the breeding conditions were the same as the above donors. Freshly ovulated eggs and milts were collected from broodstock, and after the artificial fertilization, fertilized eggs were subjected to cold shock treatment to obtain triploid. Then, treated embryos were transferred to a 1.7m^3^ seed production tank containing 15-18 °C seawater for transplantation. Hatchling larvae were reared in the microalgae environment, feeding on the intensified rotifers and brine worms, and gradually transformed into commercial feed or frozen fish.

### Isolation, identification and labeling of donor cells

Donor sexually male individuals were sacrificed by anesthetic overdose with MS-222 (Sigma-Aldrich, USA). Testes were excised and carried out cryopreservation of whole testes. The cryopreserved conditions referred to *Lee et al. (2013)*. After at least 3 d for cryopreservation, the testes were thawed. Approximately 1g of testes were dissociated followed the methods described by *Bar et al. (2015)*, with a few modifications as below. The testicular tissue was finely minced and incubated in a dissociating solution containing 40 mg/mL Collagenase H (Roche, Switzerland), 33.3 mg/mL Dispase II (Roche), 5% FBS and 1% DNase I in 2mL L-15 for 2.5 h at 25°C. The resultant cell suspension was sequentially filtered through 42um filters to eliminate non-dissociated cell clumps, suspended in a discontinuous Percoll (Sigma-Aldrich) gradient of 45%, 35%, 25% and 10%, and centrifuged at 800g for 30 min at 4°C. In addition, part cell suspension was subjected to a cell viability test by the trypan blue (0.4% w/v) exclusion assay.

After density gradient centrifugation, all cell bands were harvested and observed under microscopy (Nikon, Japan). In order to confirm donor cell band, the RNA of all cell bands and the uncentrifuged cell suspension was extracted. Based on previous researches of spermatogonial stem cell markers, the stem-related genes *oct4*, *nanos2* and *plzf* were used to identify spermatogonia by qPCR, as well as the universal germ cell marker of *vasa* as positive control (*Bellaiche et al., 2014; Lacerda et al., 2019*). The qPCR primers were shown in *Table 3*, and the method referred to *Zhao et al. (2018)*.

**Table 3.**
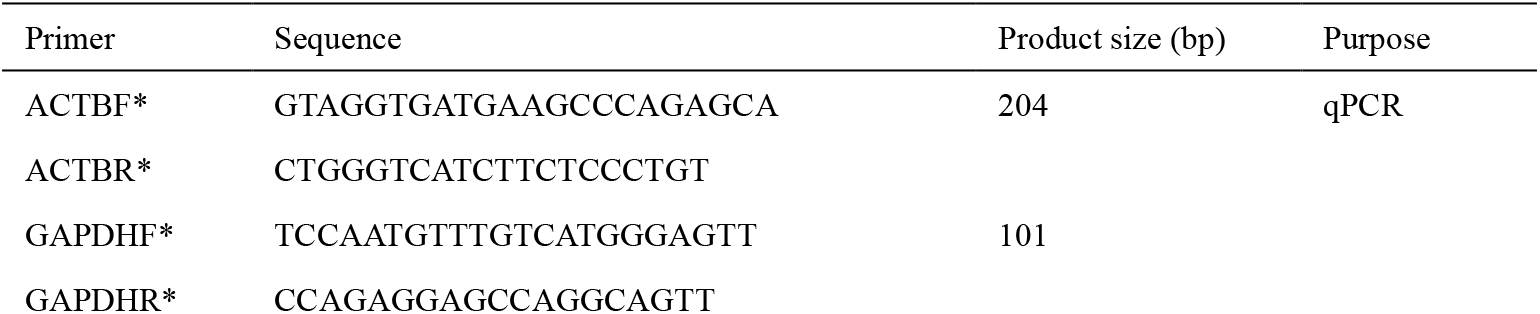
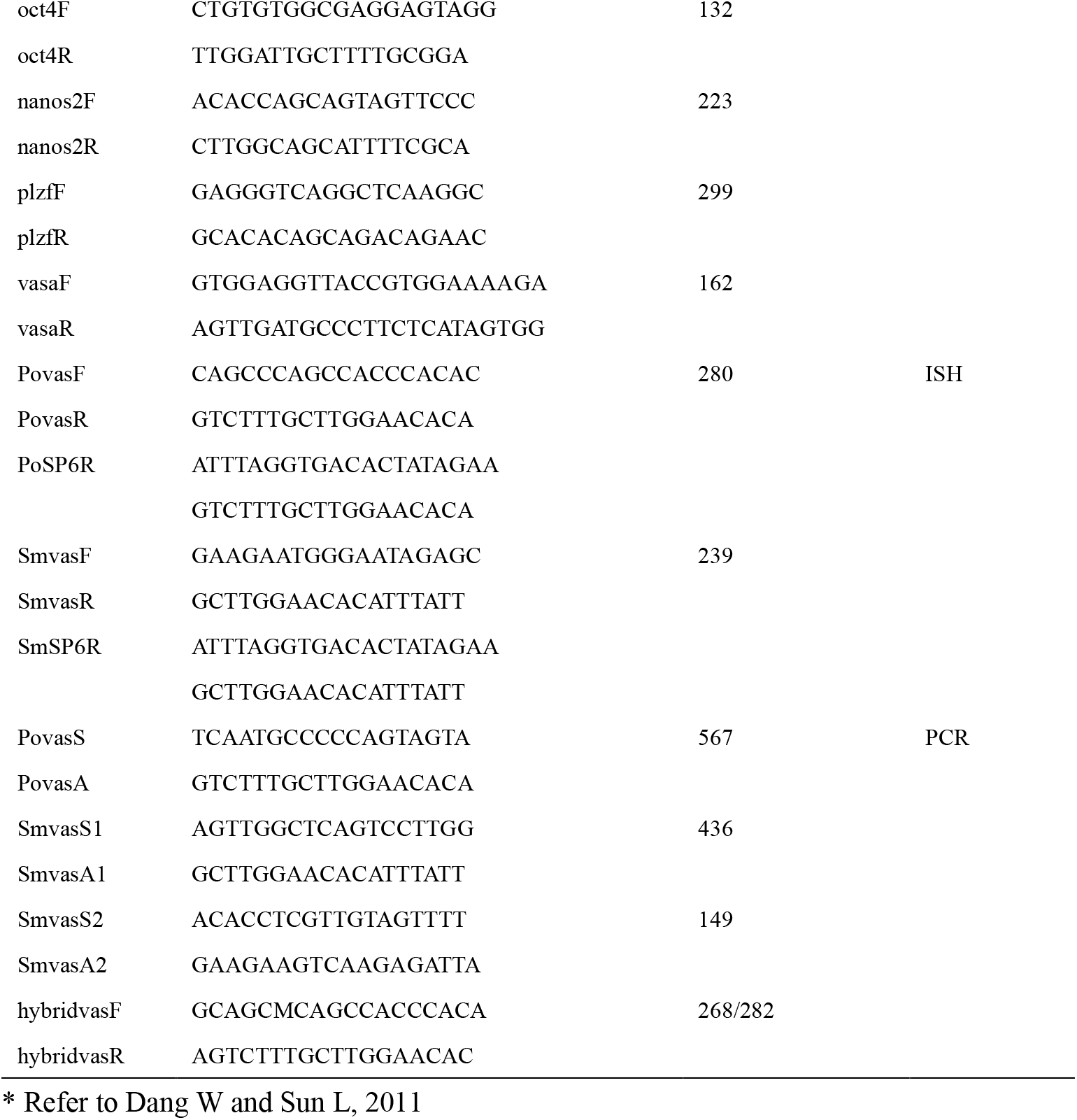
Primers used in the study

According to the identification, donor cells were recommended for optimal staining with the fluorescent membrane dye PKH26 (Sigma-Aldrich) for 5 minutes according to manufacturer’s instructions. After staining, the cells were washed twice with 3mLL-15, and resuspended in another 200ul L-15 containing 1% FBS. To detect whether the markers of PKH26 were positive cells, partial labeled cells were performed nuclear counterstain with 300 nM DAPI (Sigma-Aldrich). The labeling cells were observed and photography under fluorescent microscope (Nikon). The total number of cells was estimated, and the cells were finally diluted to 1 × 10^6^ cells / mL and kept on ice until transplantation.

### The procedures of transplantation

Japanese flounder PGCs were totally enclosed by somatic cells at 22 days post hatching (dph) and the elongated gonadal primordia appeared under the ventral kidney (*Yang et al., 2018*), so all transplantation experiments were carried out with triploid Japanese flounder before 22dph. Every type of transplantation was repeated at least two times with more than 500 larvae, as shown in *Table 1*. Transplantation needles were prepared by pulling thin glass capillaries (WPI, USA) using an electric puller (Narishige, Tokyo, Japan). The tips of the needles were ground to a 35° angle and an opening of 20-30 um diameter.

For transplantation, triploid Japanese flounder larvae were anaesthetised with 40ng/ul MS-222 in seawater, and transferred onto a Petri dish coated with 2% agar. Donor cells were then transplanted into the coelomic cavity of the anaesthetized larvae using an oil pressure manual microinjection pump (Narishige) under a stereomicroscope. After transplantation, recipient larvae were transferred from the Petri dish to a 5L recovery round plastic box filled with seawater containing 0.1% BSA and 15mg amoxicillin for three days, then transferred into nylon net hanging in the tank for two months, and released into the a 1.7m^3^ larval rearing tank finally. Non-transplanted triploid larvae were also stocked at larval rearing tanks as control.

### Fluorescent observation of donor-derived cells

Post-transplantation analysis of donor cells fate in triploid Japanese flounder recipients was first performed by fluorescent microscopy at 14 and 50 days after transplantation (dpt). To confirm that PKH26-labeled donor cells were present in the peritoneal cavity and migrated to the genital ridge, 10 recipients were randomly selected to overall observe the abdomen at 14 dpt. And the ratios of donor cells migrated to the genital ridge of recipient were counted at 14 dpt. At 50 dpt, gonads were excised or exposed by tearing abdominal skin from 5 random recipients, then washed in 1× PBS and observed with the fluorescent microscope for examination of the distribution of donor cells.

### Histology and *in situ* hybridization

All gonads were removed and fixed overnight with Bouin’s solution and 4% paraformaldehyde (PFA), respectively. Fixed samples were dehydrated through a graded series of ethanol, embedded in paraffin wax, and cut into 4um thick sections and stained with H&E.

The turbot cells could be detected in triploid Japanese flounder by *in situ* hybridization (ISH) with species-specific probes. The primers (*Table 3*) used for probe synthesis were designed in the *vasa* 3’UTR regulatory region with low homology. For the antisense RNA probe synthesis process, briefly, the synthesis templates were the introduction of SP6 at the 3’ end of the fragment by 2 rounds of PCR; RNA probes were synthesized by in vitro transcription under the drive of SP6 promoter with the DIG RNA Labeling Kit (Roche, Mannheim, Germany); Then, RNA probes were purified with SigmaSpin^™^ Sequencing Reaction Clean-Up (Sigma–Aldrich). The gonad tissues were cut into 7um thick sections, de-waxed and rehydrated. After washed with PBST, they were re-fixed using 4% PFA in PBS and digested with proteinase K (10 ug/ml) for 10 minutes. The ISH were performed with the probes at 65 °C for 14 hours and chemical stain with BCIP/NBT substrates.

### Immunohistochemistry

In order to investigate whether donor-derived cells could proliferate in the xenogenic recipients, the gonad sections of 125dpt recipient after ISH were analyzed by immunohistochemistry. The sections were blocked in normal goat serum, treated with a 1:100 diluted anti-proliferating cell nuclear antigen (anti-PCNA) antibody (KGA324, Keygen Biotech Co., China) for 1 hour, and treated with a 1:50 diluted goat anti-mouse IgG-FITC secondary antibody (KGA324). After staining, the sections were incubated with DAPI staining solution for 5 min. The sections were washed and observed through a fluorescent microscope (Nikon).

### PCR detection donor-derived spermatozoon or offspring

The spermatozoon of 5 recipients at 2 ypt with turbot donor cells were released by squeezing the abdomen of the fish. Whole genomic DNA was extracted from the spermatozoon, and PCR was performed with specific primers designed in *vasa* 3’UTR (*Table 3*). PCR reactions were carried out in 25ul volume with Q5® Hot Start High-Fidelity 2Χ Master Mix (NEB, USA), according to the routine PCR procedure of Q5. An appropriate annealing temperature was estimated by using the NEB Tm Calculator. In order to detect a small amount of turbot DNA, nested PCR was carried out with 0.5ul PCR production as template.

The spermatozoon of 3 recipients with summer flounder donor cells were collected as above, and performed artificial insemination with Japanese flounder eggs. Whole genomic DNA was extracted from 15 hybrid offspring and used for PCR to detect donor-derived DNA. The PCR was carried out with Q5, and primers designed in *vasa* 3’UTR (14bp difference between 2 species) were shown in *Table 3*. The PCR products were separated by electrophoresis on 2% agarose gel, then sent to sequence at Sangon Biotech.

### Scanning electron microscopy

Subtle morphological and structural differences of turbot and Japanese flounder could be distinguished by scanning and transmission electron microscopy. Spermatozoon were fixed in 2.5% glutaraldehyde diluted in PBS (pH 7.6), dehydrated in a series of increasing concentrations of ethanol, critical-point dried, evaporated with gold, and examined with a scanning electron microscope (JEOL JSM-840 SEM).

### Parentage test

The spermatozoon from different recipients at 2 ypt were carried out artificial insemination with turbot or Japanese flounder eggs. Meanwile, the spermatozoon (malformed) of triploid male controls were inseminated with Japanese flounder eggs. The fertilization rate and hatching rate were counted, and embryo development was tracked regularly. The offsprings of turbot and the Japanese flounder were obviously different in appearance and can be distinguished by the naked eye. Early in the late embryonic development, it could also be identified through a microscope. But the hybrids of Japanese flounder and summer flounder were only identified by the above PCR. The recipients from intra-species transplantation could evaluated by comparison with controls.

## Additional information

### Funding

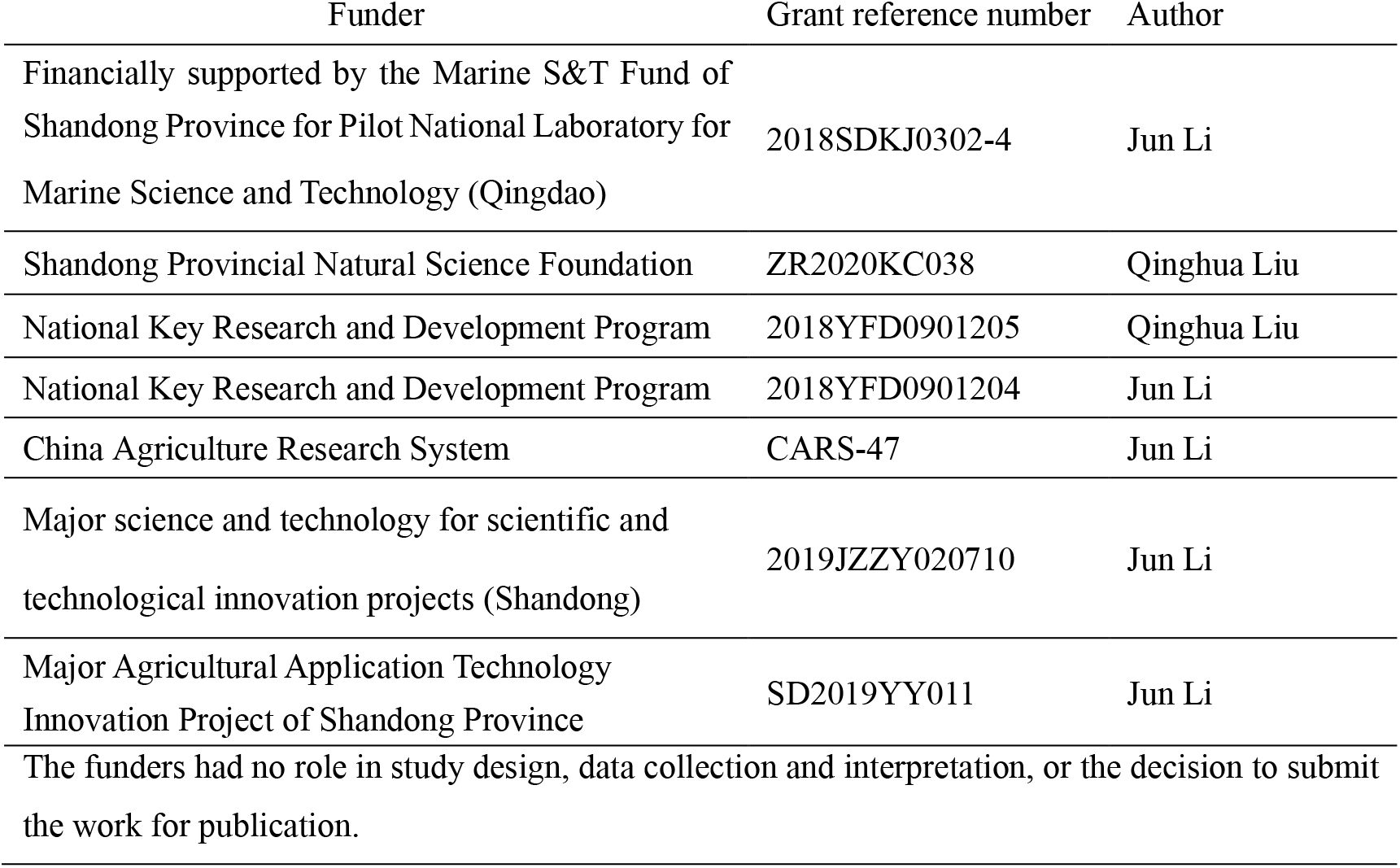

### Author contributions

Li Zhou, Conceptualization, Data curation, Formal analysis, Supervision, Validation, Investigation, Visualization, Methodology, Writing - original draft, Project administration, Writing - review and editing; Qinghua Liu, Jun Li, Investigation, Resources, Supervision, Funding acquisition, Writing - review and editing; Xueying Wang, Jingkun Yang, Conceptualization, Supervision, Investigation, Visualization, Methodology; Zhihao Wu, Feng You, Triploid Preparation; Shihong Xu, Yanfeng Wang, Zongcehng Song, Donors acquisition, Recipients cultivation.

### Ethics

All experiments were performed in accordance with the relevant national and international guidelines and approved by the Institutional Animal Care and Use Committee, Institute of Oceanology, Chinese Academy of Sciences.

